# Photoacoustic imaging resolves species-specific differences in haemoglobin concentration and oxygenation

**DOI:** 10.1101/2020.06.23.167502

**Authors:** Lina Hacker, Joanna Brunker, Ewan St. John Smith, Isabel Quiros-Gonzalez, Sarah E. Bohndiek

## Abstract

**Introduction:** Photoacoustic imaging (PAI) enables the detection of blood haemoglobin (HB) concentration and oxygenation (SO_2_) with high contrast and resolution. To date, the relationship between photoacoustically determined total Haemoglobin (THb^MSOT^) and oxygen saturation (SO_2_) biomarkers and the underlying biochemical blood parameters has yet to be established. We sought to explore these relationships in a species-specific manner.

**Methods:** Experiments were performed *in vitro* using tissue-mimicking agar phantoms. Blood was extracted from mouse, rat, human and naked mole-rat (*Heterocephalus glaber*), anticoagulated in EDTA and measured within 48 hours. THb^MSOT^ and SO_2_^MSOT^ were measured using a commercial photoacoustic tomography system (InVision 128, iThera Medical GmBH). Biochemical blood parameters such as haemoglobin concentration (HB, g/dL), haematocrit (HCT, %) and red blood cell count (RBC, μL^-1^) were assessed using a haematology analyser (Mythic 18 Vet, Woodley Equipment).

**Results:** A significant correlation was observed between THb^MSOT^ and biochemical HB, HCT and RBC in mouse and rat blood. Moreover, PAI accurately recapitulated inter-species variations in HB and HCT between mouse and rat blood and resolved differences in the oxygen dissociation curves between human, mouse and rat. With these validation data in hand, we applied PAI to studies of blood obtained from naked mole-rats and could confirm the high oxygen affinity of this species in comparison to other rodents of similar size.

**Conclusion:** In summary, our results demonstrate the high sensitivity of photoacoustically determined biomarkers towards species-specific variations *in vitro*.

## Introduction

Photoacoustic imaging (PAI) is an emerging modality able to reveal high image contrast, arising from optical absorption in tissue, at high spatial resolution, afforded by ultrasound detection. PAI is based on the photoacoustic principle^1^, whereby pulsed light is absorbed by chromophores, resulting in the generation of pressure waves that can be detected by ultrasound transducers at the tissue surface. Applying PAI at multiple wavelengths enables non-invasive, label-free detection of total haemoglobin concentration (THb) and oxygenation (SO_2_) based on the different optical absorption spectra of deoxygenated (HbR) and oxygenated haemoglobin (HbO_2_)^2^. The relative concentrations of these respective chromophores can then be calculated by spectroscopic inversion^3^. PAI measures of THb (HbR+HbO_2_) and SO_2_ (HbO_2_/THb) have been widely exploited to characterize tissues in the context of different pathologies, for example in breast cancer^4–6^, melanomas^7,8^, prostate cancer^9–11^, nodal lesions of the head and neck^12^, vascular diseases^13–15^ and colitis^16^. Particularly in cancer biology, THb and SO_2_ have proved to be of high value by allowing the detection of two cancer hallmarks: angiogenesis and hypoxia^17^.

Despite their extensive use in PAI, the relationship between photoacoustically determined THb and SO_2_ biomarkers and the underlying physiological variations in biochemical blood parameters, such as haemoglobin (HB) concentration (g/dL), haematocrit (HCT, %) and red blood cell count (RBC, μL^-1^), has yet to be established. HB is an iron-containing haem tetramer composed of two α- and two β-monomers and responsible for oxygen transport in all vertebrates^18^, yet HB-related blood parameters differ within and between species and human race^19,20^, with age and sex^21–23^ of the individual, and can change under pathophysiological conditions or pharmaceutical treatment^24^. Such differences could lead to substantial variations in PAI biomarkers or, for example, signal dynamics during more complex protocols such as a gas challenge^10^, which could lead to misinterpretation. Moreover, genetic modifications of the HB protein sequence resulting from either accumulated evolutionary changes or spontaneous mutations can alter the oxygen binding capabilities of the HB molecule^25^.

Here, we investigate the relationship of photoacoustically determined THb and SO_2_ biomarkers with biochemical blood parameters taken from mice, rats and humans in a controlled tissue-mimicking phantom system. Having examined the intra-species homogeneity of the globin genes, we first elucidate intra- and inter-species differences between photoacoustic THb and static biochemical blood parameters in mouse and rat. We then move on by examining the differences in the dynamics of the oxygen dissociation curves, here including human as a further species. With these validation data in hand, we then apply PAI to studies of blood obtained from naked mole-rats (*Heterocephalus glaber)*, a species remarkable for its resistance to oxygen deprivation^26^. Our results confirm the high sensitivity of photoacoustically determined biomarkers towards species-specific variations *in vitro*.

## Material and Methods

### 1. Sequencing

DNA was isolated from liver samples of female C57BL/6J mice (3-4 months; n=3) and Wistar rats (6-9 months; n=3) (Charles River) using the Qiagen DNeasy Blood/Tissue kit following the manufacturer’s instruction. Primers (displayed in Table 1) for both the forward and reverse strand of each gene were designed using Primer 3 software (version 4.0.0)^27^. PCR amplification was performed using the Q5® Hot Start High-Fidelity DNA Polymerase (New England BioLabs Inc.) and the corresponding protocol^28^ with an annealing temperature of 66°C. Products were purified and sequenced using Sanger sequencing (SourceBioscience). Sequences were aligned using Clustal Omega^29^. The identity match between the sequences was calculated.

**Table 1:**
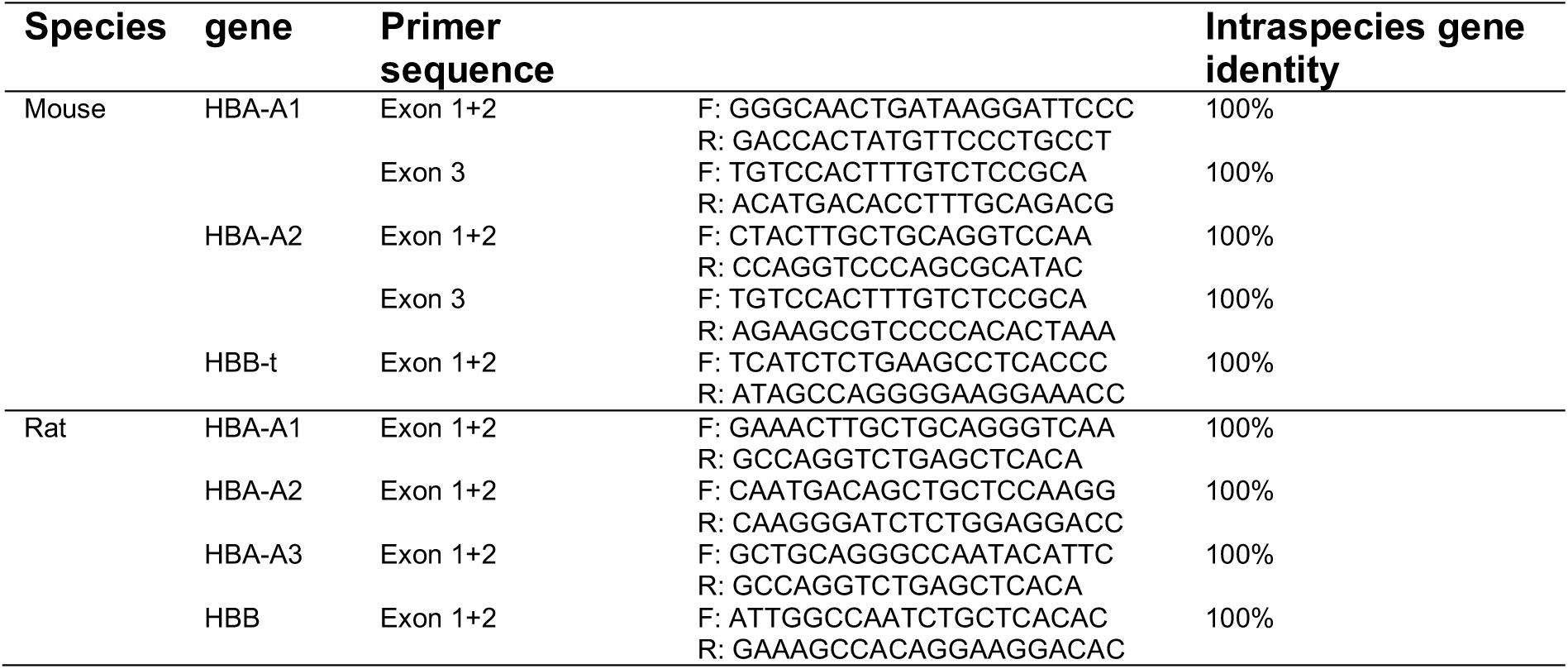
Confirmed intraspecies gene sequence identity of haemoglobin α- and β-genes in female mouse (n=3) and rat (n=3).

### 2. Blood samples

Whole blood and tissue samples (liver) for DNA extraction originating from female C57BL/6J mice (3-4 months) and Wistar rats (6-9 months) were ordered from Charles River Laboratories. For comparison of the blood parameters in section 1, 2 and 3, the same sex was chosen as HB parameters have been shown to be sensitive to the sex of individuals^22^. For the blood oxygenation experiments in the remaining sections 4 and 5, blood from mixed sex was used to achieve more generalizable results. Human blood samples from healthy donors were collected under the research ethics approval of the Royal Papworth Hospital tissue bank (project number T02196) in Cambridge. Blood from naked-mole rats (15-23 months, all male) was collected as a post mortem non-regulated procedure following decapitation of the animal for another scientific purpose. All blood samples were anticoagulated in ethylenediaminetetraacetic acid (EDTA), directly stored at 4 °C and processed within 48 hours.

### 3. Blood analysis

Blood parameters were assessed using an impedance-based haematology analyser (Mythic 18, Woodley Veterinary Diagnostics^30^). The parameters obtained included: absolute and relative number of lymphocytes, monocytes, and granulocytes; absolute number of red blood cells (RBC, μL^-1^) and white blood cells (WBC, μL^-1^); haemoglobin concentration (HB, g/dL); mean corpuscular haemoglobin (MCH, pg); mean corpuscular volume (MCV, µm^3^); mean corpuscular haemoglobin concentration (MCHC, g/dL); and haematocrit (HCT, %). MCV, MCHC and MCH are commonly defined as: MCV=[HCT]/RBC; MCHC=[HB]/[HCT] and MCH = [HB]/RBC, where square brackets denote concentrations.

### 4. Absorption spectra measurement

The absorption profile of HB was independently verified. Whole blood was lysed in distilled water (1:1) and the HB extracted in order to avoid artefacts caused by light scattering of the erythrocytes. For HB extraction^31^, blood was centrifuged three times at 3000 rpm for 3 minutes and washed with phosphate buffered saline (PBS) to remove the plasma. Afterwards, 1 unit volume of blood was thoroughly mixed with 1 unit volume of deionized water and 0.4 unit volume of toluene. The mixture was stored at 4°C for at least 24 hours to ensure complete haemolysis. The solution was then centrifuged at 13000 rpm for 10 minutes. The lowest layer containing the HB was extracted by syringe and filtered through a syringe filter (Millex, Millipore) with pore size of 0.22 µm to remove the cell debris and large particles. The absorption spectrum of extracted HB was recorded in the range of 600-900nm using a microplate reader (Clariostar, BMG Labtech).

### 5. Phantom preparation and photoacoustic imaging

Agar phantoms were produced according to the protocol by Joseph et al^32^. Briefly, liquid 1.5% w/v agar (Fluka 05039) was mixed with 2.1% v/v pre-warmed intralipid (Sigma-Aldrich I141) to provide a reduced scattering coefficient of 5cm^-1^. Nigrosin dye (Sigma-Aldrich 198285) was added to provide an absorption coefficient of 0.05cm^-1^ (at 564 nm, peak of the nigrosin spectrum). The solution was poured into a 20mL (2cm diameter) syringe with the injection end removed and with polyvinyl chloride (PVC) tubing (inner diameter: 1.5mm, outer diameter: 2.1 mm; VWR 228-3857) inserted along the central axis. After the agar was set, the phantom was removed from the syringe ready for imaging.

PAI experiments were performed using a commercial small animal imaging system (MSOT inVision 256-TF; iThera Medical GmbH). The system has been described in detail elsewhere^33,34^. Briefly, a tunable (660–1300 nm) optical parametric oscillator (OPO), pumped by a nanosecond (ns) pulsed Nd:YAG laser, with 10Hz repetition rate and up to 7ns pulse duration provides excitation pulses. Light is delivered to the sample through a custom optical fibre assembly, which creates a diffuse ring of uniform illumination over the imaging plane. The sample is coupled to the transducers by using a water bath, filled with degassed and deionized water. For ultrasound detection, 256 toroidally focused ultrasound transducers covering an angle of 270° are used (centre frequency of 5MHz, 60% bandwidth) allowing tomographic reconstruction. A minimum of four images were taken along the length of the phantom at steps of 0.5mm using seven wavelengths (700nm, 730nm, 760nm, 800nm, 850nm, 900nm, 1040nm) with an average of 10 pulses per wavelength.

For the measurements of the oxygen-disassociation curve (ODC), a flow phantom setup was used, which has also been described in detail elsewhere^35^. For each measurement, 5mL of pooled blood from the respective species was first oxygenated by the addition of 0.2 % v/v hydrogen peroxide (H_2_O_2_ 30% (w/w) in deionized water, Sigma-Aldrich 7722-84-1). The oxygenated blood was filled into the circuit, carefully avoiding the introduction of air bubbles. During the course of the experiment, a syringe driver (Harvard, MKCB2159V) was used to deoxygenate the blood with 0.03 % w/v sodium hydrosulfite (ACROS Organics 7775-14-6) in PBS) at a constant flow rate of 10 µL/min. The experiment was performed at room temperature and a peristaltic pump (Fisher Scientific CTP100) was used for blood circulation. Oxygen fluorescence quenching needle probes (Oxford Optronix, NX-BF/O/E) were placed before and after the tissue-mimicking phantom, which recorded the temperature and partial pressure of oxygen (pO_2_, mmHg) in real time. The data were downloaded via an Arduino UNO and read in MATLAB. Using the same commercial PAI system as above, images were acquired at a single position (no pulse-to-pulse averaging) for 17 wavelengths (660 nm, 664 nm, 680 nm, 684 nm, 694 nm, 700 nm, 708 nm, 715 nm, 730 nm, 735 nm, 760 nm, 770 nm, 775 nm, 779 nm, 800 nm, 850 nm, 950 nm). Absorption spectra were measured using a light source (Avantes Avalight-HAL-S-Mini) and spectrometer (AvaSpec-ULS2048-USB2-VA-50). Absorption spectra were recorded continuously via AvaSoft software as the fluid passed through a flow cell (Hellma Analytics, 170700-0.5-40) as it has been shown that directly measured absorption spectra provide the most accurate endmembers for spectral unmixing^35^. Species-specific absorption spectra at the point of complete oxygenation and deoxygenation were extracted and used for spectral unmixing of the data recorded for the respective species. Between each measurement run the tubing containing the blood was cleaned with phosphate buffered saline (PBS).

For the experiments involving naked mole-rat blood a simpler setup was used as only a limited amount of naked mole-rat blood could be obtained. Blood samples (100µL) were inserted into a straw within a phantom and 10 μL H_2_O_2_ (0.03% in PBS) was injected using a syringe. The oxygenation of the blood was directly measured after injection using MSOT. 30 images were taken of a chosen slice using 7 wavelengths between 660 nm and 1140 nm, with an average of 10 pulses per wavelength. Each image took 8s to acquire. Measurements were conducted at 37**°**C. It should be noted here that naked-mole rats are considered poikilothermic and usually have a physiological body temperature of around 30-32**°**C^36^. Examining the oxygen affinity at higher temperature decreases the oxygen affinity of the blood^37^. However, it has been shown that even at 37**°**C significant differences between the oxygen affinity of naked mole-rat and mouse are still present^38^, supporting our experimental approach.

### 6. Image and statistical analysis

PAI data were analysed using the ViewMSOT software (v3.6.0.119; iThera Medical GmbH). Model–based image reconstruction and linear multispectral processing were applied to extract the relative signal contributions of HbO_2_ and HbR. The same position within the phantom was used to determine the average intensities for HbO_2_ and HbR. Regions of interest (ROIs) were manually drawn around the circular cross section of the tube inserted in the phantom. THb^MSOT^ was calculated as the sum of HbO_2_ and HbR. SO_2_^MSOT^ was calculated as the ratio of HbO_2_ to THb^MSOT^ signal. The expressions ‘THb^MSOT^’ and ‘SO_2_^MSOT’^ are used to emphasise that the photoacoustically determined THb and SO_2_ biomarkers are not exactly equal to the underlying parameters, since to be able to accurately resolve absolute values, knowledge of the light fluence distribution, system response and Grüneisen parameter is required^39^.

Raw data extracted from the ROIs were analysed using Python (version 2.7) and MATLAB. Statistical analysis was performed using Prism (GraphPad). All data are shown as mean ± standard error of the mean (SEM) unless otherwise stated. Unpaired 2-tailed t-tests were performed to calculate the statistics. Pearson’s rank test was performed to assess correlations between biochemical blood parameters and THb^MSOT^. Significance is assigned for p-values <0.05.

## Results

### Gene sequence analysis confirms intra- and inter-species homogeneity of haemoglobin genes in mouse and rat

We first assessed the intra-species genetic correspondence of the two α- and two β-globin chains of the HB tetramer in rat and mouse, as alterations in HB globin genes can lead to changes in the protein structure that might affect the PAI signal. We found that an intra-species homogeneity of 100% could be determined (n=3, Table 1). These results indicate that intra-species signal variations in the following experiment are unlikely to be caused by genetic missense mutations, but rather by other physiological or technical sources.

### PAI THb correlates directly with biochemical HB values obtained using a haematology analyser and detects intra-species variations

The relationship between the THb^MSOT^ biomarker and biochemical blood parameters was examined. THb^MSOT^ values from photoacoustic images taken from our static blood phantoms correlated with biochemical HB (Figure 1A), HCT (Figure 1B) and RBC (Figure 1C) in the same mouse blood sample (HB: Pearson r=0.6867 p=0.0047; HCT: Pearson r=0.6083, p=0.0161; RBC: Pearson r=0.5594, p=0.0302). To determine whether these observations are species-independent, the correlation between THb^MSOT^ and biochemical blood parameters was also studied in rat. Again, a significant correlation was found between THb^MSOT^ and biochemical HB (Figure 1D), HCT (Figure 1E) and RBC (Figure 1F) in rat blood (HB: Pearson r=0.6786, p=0.0047; HCT: Pearson r=0.5474, p=0.0427; RBC: Pearson r=0.5363, p=0.0480) suggesting that PAI is directly sensitive to intra-species variations in HB, HCT and RBC parameters.

**Figure 1:**
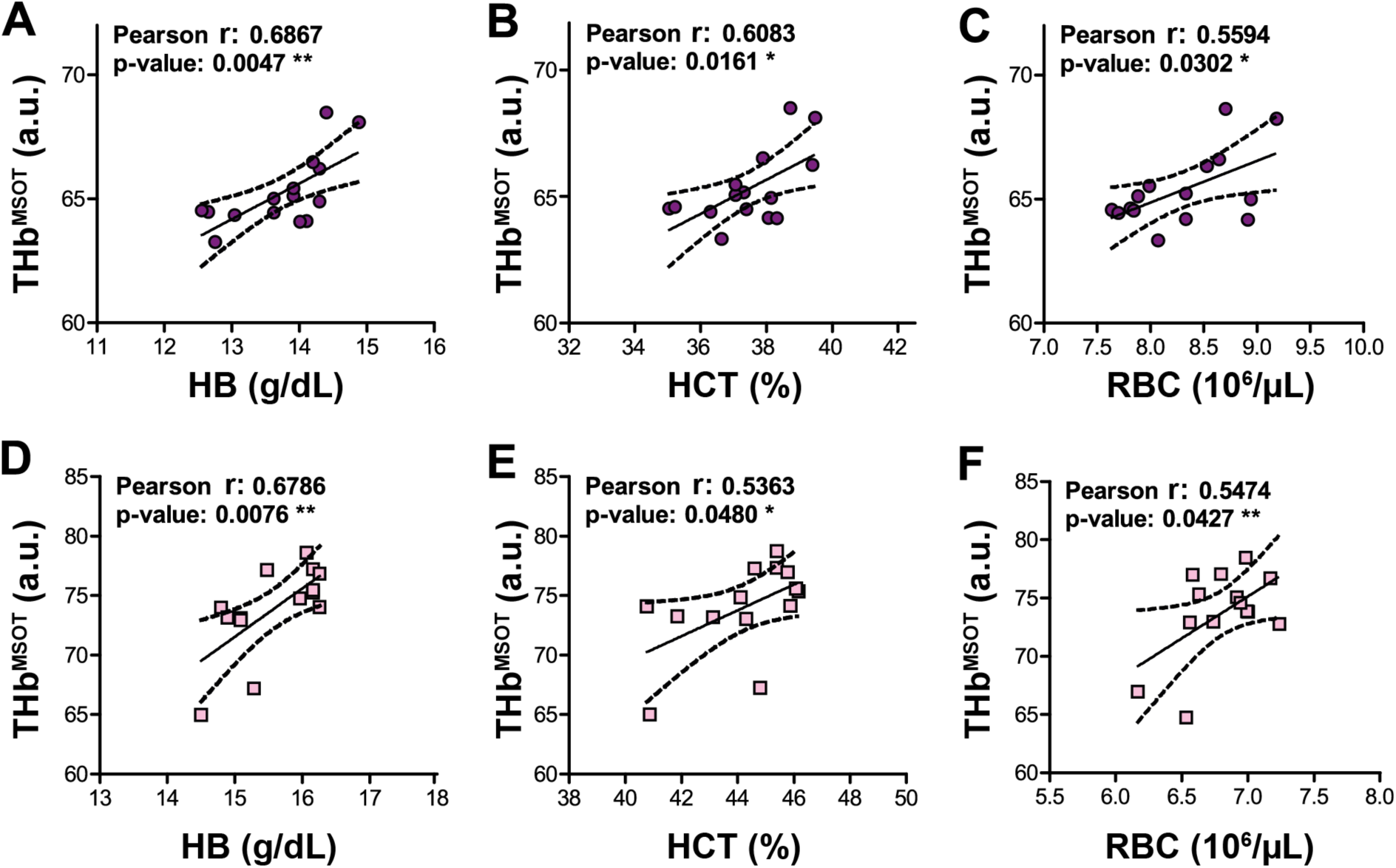
Total haemoglobin THb^MSOT^ biomarker measured in tissue-mimicking phantoms corresponds to biochemical HB, HCT and RBC levels in mouse and rat. THb^MSOT^ extracted from PAI data obtained from tissue-mimicking phantoms containing blood from female mice (n=15, purple circles, panels A - C) and rats (n=14, pink squares, panels D - F) correlated significantly with biochemical HB (A,D), haematocrit HCT (B,E) and red blood cell count RBC (C,F) concentration in both species. * p<0.05, ** p<0.01 by Pearson correlation.

### PAI THb resolves inter-species differences in HB and HCT *in vitro*

After assessing the impact of intra-species variation in biochemical blood parameters on MSOT signal, the effect of inter-species differences was analysed. In order to exclude major differences in the protein structure and conformation yielding differences in the absorption spectra for the haemoglobin, we first confirmed that no inter-species difference in the absorption spectra of HB extracted from mouse and rat blood was detected. Minor differences in total light absorbance of the samples were observed, which could be explained by different HB levels of rat and mouse blood (Table 2); thus, absorption spectra were normalized to the maximum absorption at 450 nm for further comparison. In line with the literature^40–42^, the experimentally determined spectra of mouse and rat blood demonstrated a very good agreement (Figure 2A).

**Table 2:** Comparison of experimental and literature HB values of mouse and rat.

| Parameter | Mouse (BALB/c) | Rat (Wistar) |
| --- | --- | --- |
| HB (exp.) (g/dl) (mean±std) | 14.73±2.09 | 15.06±2.35 |
| HB (lit.) (g/dl) | 12-17 | 14.4-18.0 |
| HCT (exp.) (%) (mean±std) | 41.83±6.38 | 45.50±6.13 |
| HCT (lit.) (%) | 40-68 | 36-48 |
| Reference | James Fox et al<br>2006<sup>61</sup> | Charles River Laboratories, 1998<sup>62</sup> |

**Figure 2:**
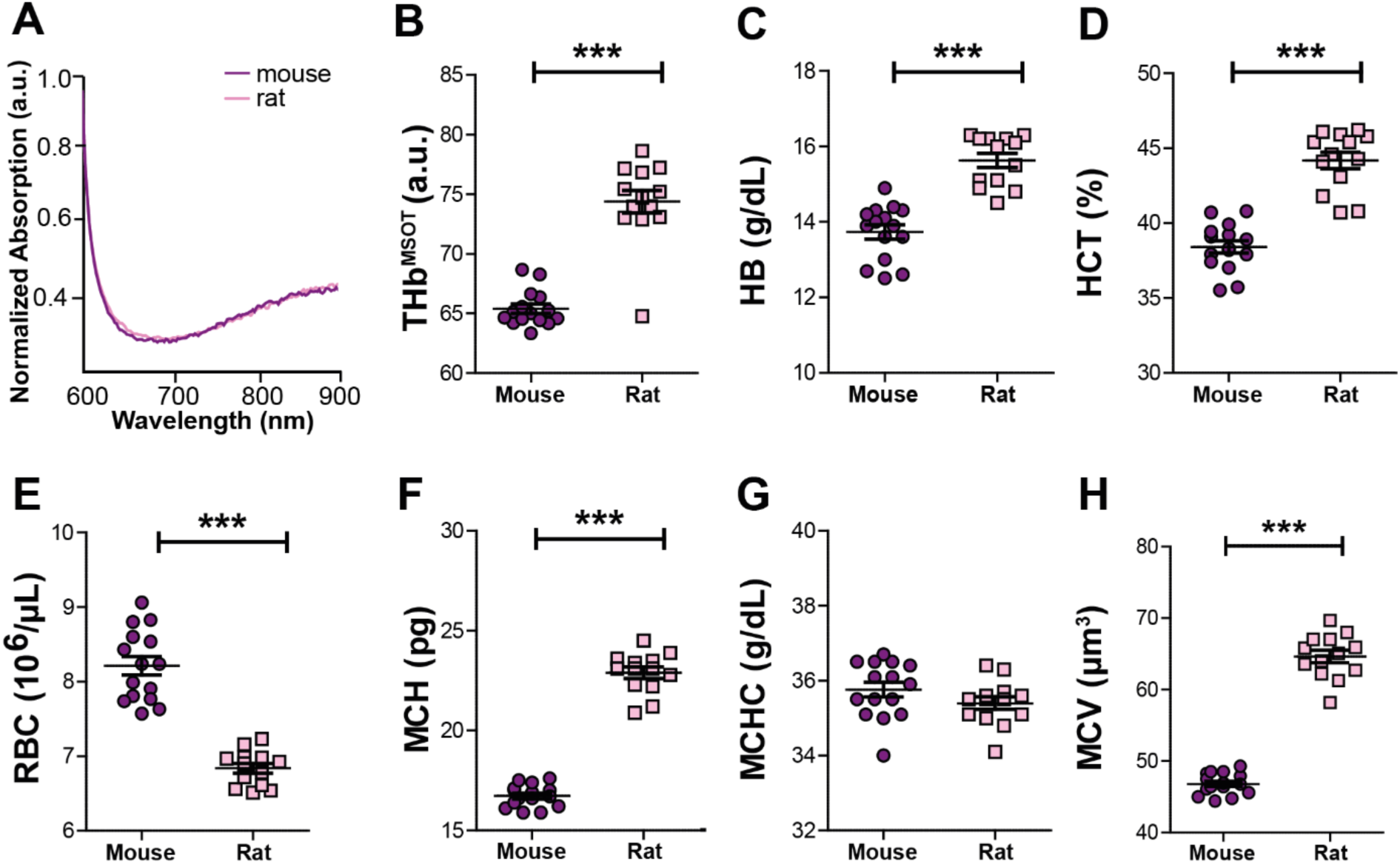
PAI resolves inter-species HB differences in rat and mouse. **(A)** Absorption spectra of haemoglobin extracted from lysed whole blood of mouse (purple) and rat (pink). (B) Inter-species differences in THb^MSOT^ obtained from PAI measurements of female mouse (n=15) and rat (n=14) blood show a significantly higher value in rat blood, which was underscored by differences in biochemical HB (C), HCT (D), RBC (E), and mean corpuscular haemoglobin MCH (F). Notably, there was no difference in mean corpuscular haemoglobin concentration MCHC (G) although the mean corpuscular volume MCV (H) was also higher in the rat. Data are represented as mean ±SEM, significance *** p < 0.0001 by unpaired t-test.

Next, THb^MSOT^ was compared to the biochemical blood parameters of the different species. A significantly higher THb^MSOT^ level was observed for the rat (p<0.0001, Figure 2B). Corresponding to this observation, significantly higher HB (Figure 2C) and HCT (Figure 2D) values could be observed in this species (p<0.0001). Interestingly, the rat was characterized by significantly lower RBC levels (Figure 2E). We compared the MCH levels between the two species to establish whether this observation was associated with differences in the average HB amount per RBC. In correspondence to the HB and HCT values, significant higher MCH levels were found in the rat (Figure 2F, p<0.0001). A more detailed analysis revealed that the higher MCH values in the rat do not originate from a higher MCHC value (Figure 2G), but rather from a larger MCV of the RBC (p<0.0001, Figure 2H). These results suggest that PAI correctly resolves interspecies differences in HB and HCT *in vitro* but can only be used to quantitatively compare interspecies RBC values when MCH values are within the same range.

### PAI resolves inter-species differences in oxygenation dynamics

We next examined whether PAI has sufficient sensitivity to capture inter-species differences in oxygenation dynamics in mice, rat and human blood based on the known differences in oxygen dissociation curves (ODCs) in the literature between these species(Figure 3A)^43^. Under standard conditions (pH = 7.4, pCO_2_ = 40 mmHg (5.3 kPa), temperature = 37 °C, carboxyhaemoglobin < 2 %), human HB is known to have the highest oxygen affinity with an ODC shifted farthest to the left (p50_std_ = 26 mmHg), followed by rat (p50_std_ = 32 mmHg) and then mouse HB (p50_std_ = 48.5 mmHg)^43^ (Figure 3A).

**Figure 3:**
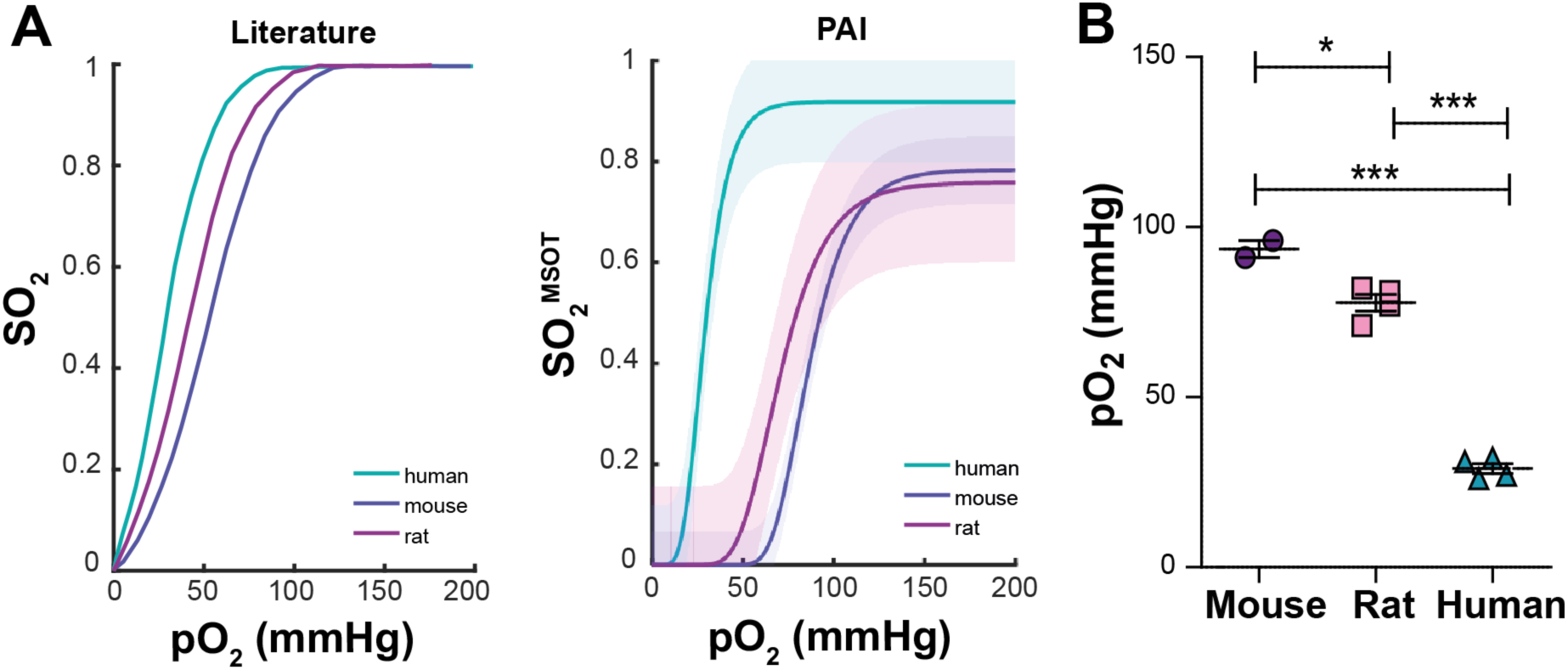
PAI resolves inter-species differences in oxygen dissociation curves (ODCs) between mouse, rat and human. Oxygenated mouse (n=2), rat (n=4), and human (n=5) blood samples were deoxygenated in a dynamic flow phantom circuit at room temperature and changes in absorption spectrum, pO_2_ and PAI signals were recorded. (A) Literature values for ODCs of the respective species under standard conditions.^43^. (B) SO_2_^MSOT^ was calculated from PAI images using the absorption spectra measured within the flow phantom circuit for spectral unmixing. The resulting values were plotted against the measured pO_2_ of the blood within the circuit at the same time point in order to produce an ODC. (C) Extracted p50^MSOT^ values denoting the pO_2_ at 50% SO_2_ ^MSOT^ are shown. It should be noted that the ODC influencing factors 2,3-DPG concentration, acid-base balance, and amount of dyshemoglobins were not standardised, but rather reflect the physiological values found in the respective species.

To test whether MSOT is able to qualitatively resolve these differences, fully oxygenated blood was introduced into a dynamic flow phantom system and the blood was gradually deoxygenated while the absorption spectra, pO_2_, temperature and PAI spectral data were recorded in real-time. The species-specific ODCs were compared by plotting sO_2_^MSOT^ against pO_2_ and showed broadly similar results as reported in the literature^25^ (Figure 3B). In similar pattern, the human ODC was found to be shifted farthest to the left (p50_MSOT_ = 29.00 ± 2.94 mmHg), followed by the rat (p50_MSOT_ = 77.75 ± 4.99 mmHg) and then the mouse (p50_MSOT_ = 93.50 ± 3.54 mmHg) (Figure 3B). These results indicate that MSOT is capable of qualitatively resolving interspecies oxygenation dynamics *in vitro*.

### PAI is sensitive to the enhanced oxygen dissociation curve of the naked mole-rat

Having established the capacity for PAI to resolve differences in both HB parameters and oxygenation dynamics in mice and rats, we made a first PAI study of blood from the naked mole-rat. The naked mole-rat is adapted to live an hypoxic underground environment, which involves having a higher affinity for oxygen in its haemoglobin molecules than rodents of similar size^38^.

As only small amounts of naked mole-rat blood could be obtained, a simpler experimental design was used for the studies in which small blood samples were oxygenated with H_2_O_2_ (0.03% in PBS) and the oxygenation measured using PAI during four minutes. A significantly higher maximum SO_2_^MSOT^ after the four minutes could be found for the naked mole-rat blood in comparison to the same experiment conducted with mouse and rat blood (Figure 4A). Considering the sigmoidal shape of the ODC, this should lead to the least inclined slope of the oxygenation change, which was indeed found in our experiments (Figure 4B). The results suggest that PAI is able to resolve the higher oxygen affinity of the naked-mole rat blood^38^ compared to mouse and rat, indicated by the overall higher SO_2_ and lower change of SO_2_ when oxygenating the samples under similar experimental conditions.

**Figure 4:**
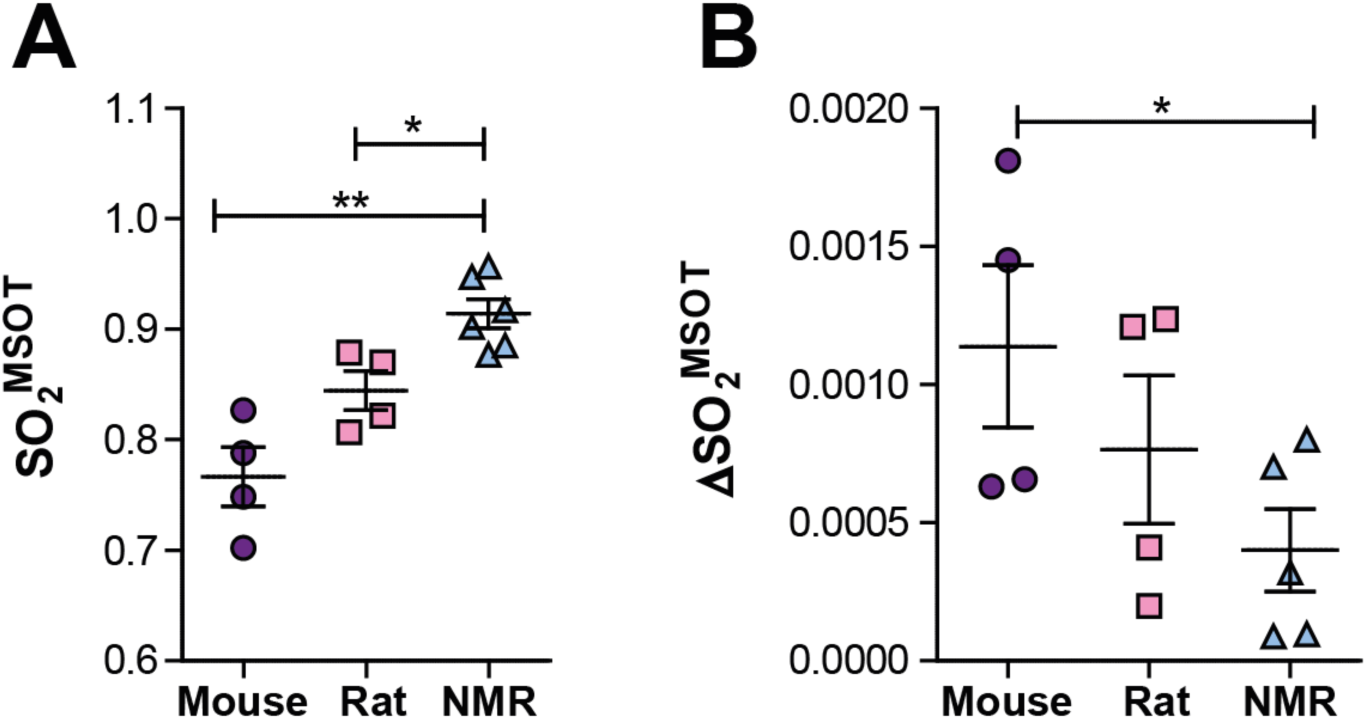
PAI evaluation of oxygenation dynamics is sensitive to the higher oxygen affinity of naked mole-rat blood. Blood samples taken from mouse (n=4), rat (n=4) and naked mole-rat (NMR; n=5) were oxygenated using 10 μL hydrogen peroxide (0.03% in PBS) at 37 ° C. (A) The maximum SO_2_^MSOT^ was highest in the naked mole-rat. (B) Considering the sigmoidal shape of the ODC, this would be expected to lead to the least inclined slope of the oxygenation change, ΔSO_2_^MSOT^, which could be confirmed in our experiments.

## Discussion

PAI holds substantial potential for application in the measurement of THb and SO_2_ biomarkers, however, our understanding of the relationship between these biomarkers and the biochemical parameters of blood including haemoglobin concentration, haematocrit, and oxygen dissociation curves, has yet to be fully elucidated. Here, we sought to study the sensitivity of PAI towards physiological variations of HB, including both intra- and inter-species variations.

Our results indicate that PAI determined THb^MSOT^ and SO_2_^MSOT^ can resolve physiological variations in HB and HCT both within and between species. We found significant linear correlations of THb^MSOT^ with HB, HCT and RBC count. As HCT and HB linearly depend on each other (roughly HB = HCT/3)^44^, it is unsurprising that both show linear trends. Our findings indicate that when the concentration of corpuscular HB is comparable between subjects, differences in RBC count could in principle be determined using PAI, however, this would require a low variance of the corpuscular HB value within the investigated group of subjects. Even within species, significant variations in MCH can occur due to disease-related macro- or microcytic anaemia or age^45^, which should be considered when making any conclusions from PAI signal to RBC number.

We could further show that differences in inter-species oxygenation dynamics based on the measurement of the PAI biomarker SO_2_^MSOT^ could be clearly resolved. This is notable given the diversity of species studied. In particular, our studies revealed that SO_2_^MSOT^ is sensitive to the differences in oxygen dissociation curves between mouse, rat and human, as well as naked mole-rat. It has to be noted that our study only aimed for a qualitative comparison of the ODC curves between the species. For a quantitative comparison of ODC curve characteristics and p50 values, other ODC influencing factors, such as species-specific 2,3-diphosphoglycerate (2,3-DPG) concentration^46^, pH^47^, amount of dyshaemoglobins^48^ or the partial pressure of carbon dioxide (pCO_2_)^49^ would also need to be taken into consideration.

Our study indicates that the application of PAI to detect functional differences in HB should be considered carefully when comparing data obtained from different species, particularly when using an “oxygen-enhanced” or “gas challenge” imaging protocol^10,50^. It also highlights the exciting potential of PAI derived biomarkers to be applied for studies in disease-associated anaemia^51^ or haemoglobinopathies, such as sickle cell anaemia^52^. Patients with haemoglobinopathies where globin proteins are structurally abnormal often show shifts in the ODC^53^, which may be captured using PAI.

Despite these promising findings, there remain some limitations of the study. We used a standard linear spectral unmixing method for resolving the contributions of HbR and HbO_2_ to our signals, which did produce values of SO_2_^MSOT^ with limited dynamic range, particularly at the higher end of the oxygen dissociation curves, when compared to ground-truth. Employing more advanced multispectral processing techniques should further enhance the accuracy of the photoacoustically determined SO_2_ estimation.

Further, our *in vitro* phantom experiments provide a well-controlled reference measurement for both the haemoglobin absorption spectrum and the partial pressure of oxygen. While it may be possible in future work to use the knowledge of inter-species variations observed in this study to make a qualitative *in vivo* comparison of the PAI biomarker response, for further validation of our findings, *in vivo* confirmation would be advantageous. Unfortunately, the experimental design would be complex because of the imaging artefacts that arise due to: tissue heterogeneities, which lead to local variations in optical and acoustic properties, and consequently to uncertainty in PAI fluence distributions^54^; motion, such as breathing or heartbeart^55^; and anatomical positioning within a non-rigid animal holder^32^. Furthermore, factors influencing blood extraction, such as blood clotting, haemolysis, dilution of blood with interstitial fluid or varying lag time before the analysis of the sample can affect the determination of the biochemical blood parameters^56^. Nonetheless, such studies are of particular importance when PAI is performed in a clinical environment, as globin gene mutations in humans are common, affecting around 7% of the overall population^37,57^. Moreover, HB concentrations have been found to vary with human ethnic group^58–60^, which could impact the acquired results in studies of mixed populations.

In summary, our findings highlight the encouraging capacity of PAI to resolve intra- and inter-species differences in HB-related blood parameters and oxygenation dynamics in a sensitive and label-free manner. These results suggest a need for careful considerations of future study designs comparing data between species, but suggest promising future avenues for application of PAI for HB- and oxygenation-related research studies, strengthening the position of PAI as a powerful and versatile tool in biomedicine.

## Acknowledgements

The authors thank: the Biorepository Unit (BRU, Matthew Clayton) of the CRUK Cambridge Institute for their assistance with mouse and rat blood; and Dr Doris Rassl and the Royal Papworth Hospital tissue bank for providing the human blood. This work was funded by Cancer Research UK under grant numbers C14303/A17197, C9545/A29580, C47594/A16267 and C197/A16465 (SEB) as well as C56829/A22053 (EStJS). LH is funded by a studentship from the National Physical Laboratory.

